# Redesign of an Undergraduate General Microbiology Lab to Include Authentic Discovery-Driven Research on Cucumber Fermentations

**DOI:** 10.1101/356261

**Authors:** Jennifer Greenstein, Brian Ford, Sandra Gove, Fred Breidt, Alice Lee, Michael E. Taveirne

## Abstract

Many undergraduate introductory microbiology laboratory courses teach basic principles of bacteriology using classical protocol-based experiments, with limited critical thinking and inquiry-based learning practices. We initiated a comprehensive redesign in our General Microbiology Laboratory course to promote scientific critical and creative thinking, while strengthening core microbiology concepts and skills. As part of the redesign, a series of authentic discovery-driven labs, based on cucumber fermentations, were developed as an independent research module within the course curriculum. Integrating discovery-driven labs allowed students to be engaged problem solvers, applying the scientific process to develop hypotheses, design experiments, utilize quantitative reasoning, and effectively communicate results. The inquiry-guided research project was developed to evaluate the minimum concentration of salt (NaCl) required in fermentation brine to safely, and effectively, ferment cucumbers. Over 5 weeks, students assess different aspects of the fermentation process, including quantifying bacterial populations with differential and selective media, measuring pH and glucose concentration of brine solutions, and characterizing the microbial metabolic potential. Additionally, students isolate an unknown bacterium from their fermentations, identifying and characterizing the isolate using 16S rRNA gene sequencing and metabolic tests. Throughout the research project, students collect, graph, and analyze their observations, culminating in students creating and presenting a scientific research poster. With this lab redesign, students generate new knowledge contributing to our understanding of microbial ecology within food fermentations, learn core microbiology skills and techniques, and develop critical and creative thinking skills. The impact of their research is valuable to science educators, researchers, and industry partners.

## INTRODUCTION

General Microbiology Lab at NC State University (NCSU) serves as an introductory microbiology lab experience for upper-level undergraduate students. Instruction in basic microbiological laboratory skills including aseptic technique, bacterial isolation (T-streak and dilutions), biochemical and physiological characterization of bacteria, and staining and light microscopy are covered. Before this lab redesign, these skills and concepts were taught using traditional protocol-based experiments with limited inquiry or critical and creative thinking, using the classic identifying an unknown bacterium project^1^. A major goal of this course redesign was to actively engage students in the creative process of scientific inquiry by incorporating authentic research into the curriculum. This type of learning environment more closely resembles a modern research lab and creates an excitement for scientific inquiry, striving to keep students in their academic trajectories as STEM majors, as well as preparing them for future scientific careers. Furthermore, this redesign aims to increase student understanding of core content, as well as introduce current microbiological and molecular techniques used in the field.

The opportunity to learn microbiological techniques and the scientific method in the context of authentic research has been shown to increase student motivation and learning gains over traditional exercises that teach or demonstrate well-documented phenomena^2-4^. Recent work has shown that students in microbiology courses have misconceptions regarding core microbiological concepts, including evolution, cell structure and function, metabolism, and genetics^5^. Furthermore, it is hypothesized that students do not remember new information simply by memorization, but through incorporating new information in context with previous knowledge^6^. Course-based Undergraduate Research Experiences (CUREs) can aid in increasing student understanding of core content as well as increase critical and creative thinking skills^7-9^. In this course redesign, students rely upon their abilities to think critically and creatively to generate questions, develop hypotheses, design experiments, record and analyze observations, draw conclusions, and propose future directions as part of an authentic research project. To accomplish these goals, this course features a series of interconnected experiments in three distinct modules, focused on the science of microbial food fermentations. Learning the principles of food microbiology offers an opportunity to teach basic microbiological concepts with a subject relatable to students. Topics including food safety, foodborne infections, public health, and microbial industrial practices are relevant concepts that are discussed in context of this research project.

Many of the foods we eat are products of microbial processes. Fermentation is a fundamental microbial process used in making yogurt, cheese, and pickled vegetables. In fruit and vegetable fermentations, salt brines are used to limit the growth of spoilage organisms (i.e. *Enterobacteriaceae*) while permitting the growth of beneficial lactic acid producing bacteria, (i.e. *Lactobacillaceae*)^10,11^. Growth of members of the *Lactobacillaceae* family result in organic acid production, specifically lactic acid, which is an inhibitory environmental factor preventing the growth of spoilage microorganisms in addition to the salt brine. Although salt brines aid in safe and effective fermentations, the resulting waste contains high concentrations of salt, specifically sodium chloride (NaCl), that must be treated prior to disposal^12^. The authentic research project in this course redesign studies the succession of bacterial communities of cucumber fermentations in different salt concentrations to identify the lowest NaCl concentration in brine solutions needed to create a safe, and commercial food product. Throughout the semester, students work in pairs, and groups, to assess the progress of an experimental cucumber fermentation by collecting, graphing, and interpreting data, culminating in a final electronic poster presentation of their work.

### Intended Audience

General Microbiology Lab is an introductory, large-enrollment, critical-path course enrolling ~650 upper-level (junior and senior) undergraduate students per year. Each semester there are 14 sections (~24 students/section) taught by Graduate Teaching Assistants, supervised and mentored by a Teaching Assistant Professor. Prerequisites for this course include one semester of introductory biology and one semester of organic chemistry. A General Microbiology lecture course is a separate, required, co-requisite for the lab. Most students registered for the laboratory course at NCSU are life science majors.

### Learning Time and Learning Objectives

General Microbiology Lab is a 12-week, one-credit hour course, where students meet once a week for 2 hours 45 minutes. An approximately 30-minute lecture discussing lab procedures and theory precedes the laboratory experiments. The redesigned course is divided into three distinct modules. Module 1, taught in the first four weeks of lab, introduces students to core skills in microbiology. Module 2, which encompasses the inquiry-guided research project, is carried out over five separate lab sessions. Module 3, which covers data visualization and presentation, is taught over two individual lab sessions. The last lab is the final exam and lab practical assessing microbiology techniques and content. The learning objectives for this course are mapped to the ASM Recommended Curriculum Guidelines for Undergraduate Microbiology Education^13^, highlighted in Table 1.

**TABLE 1.**
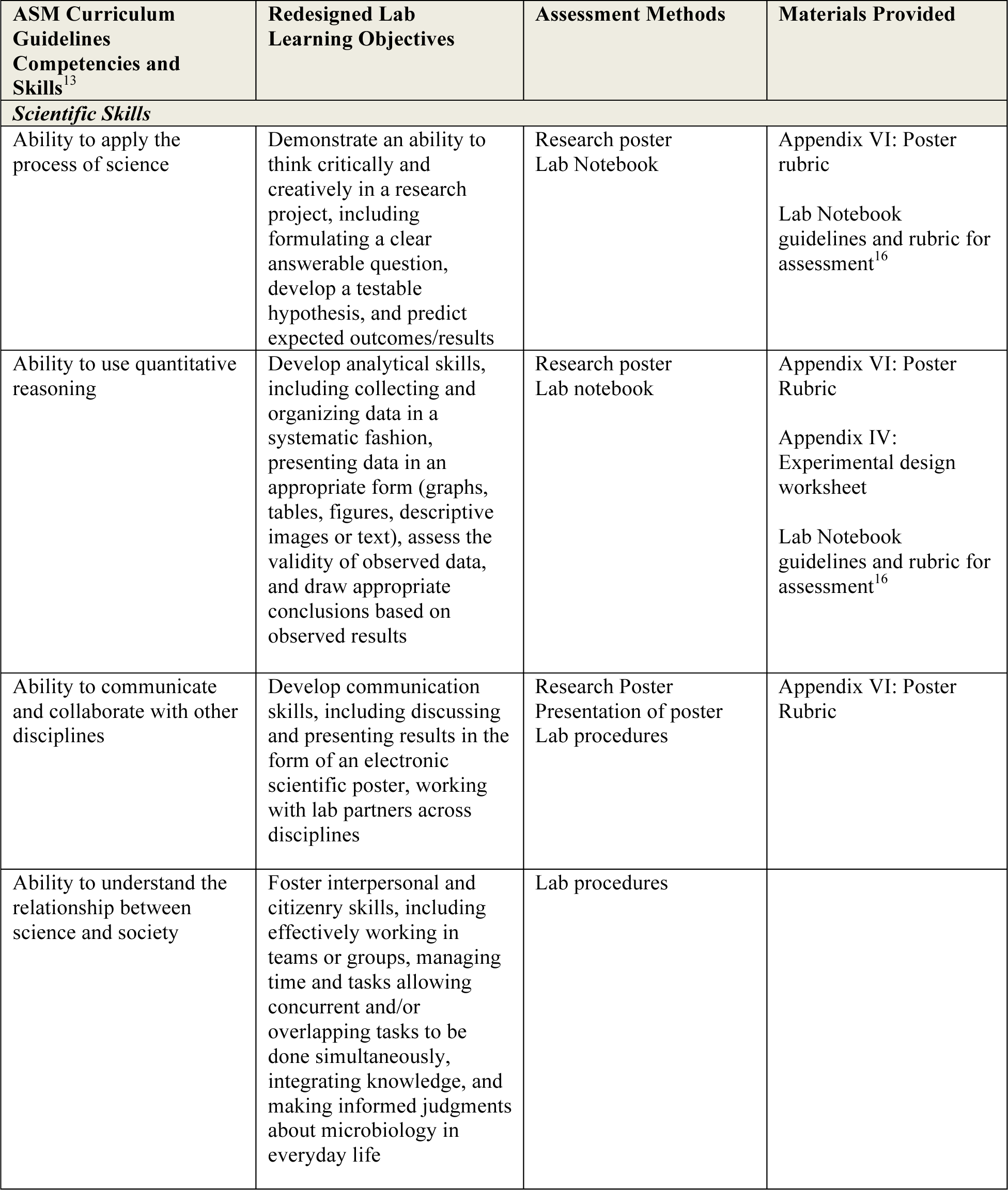
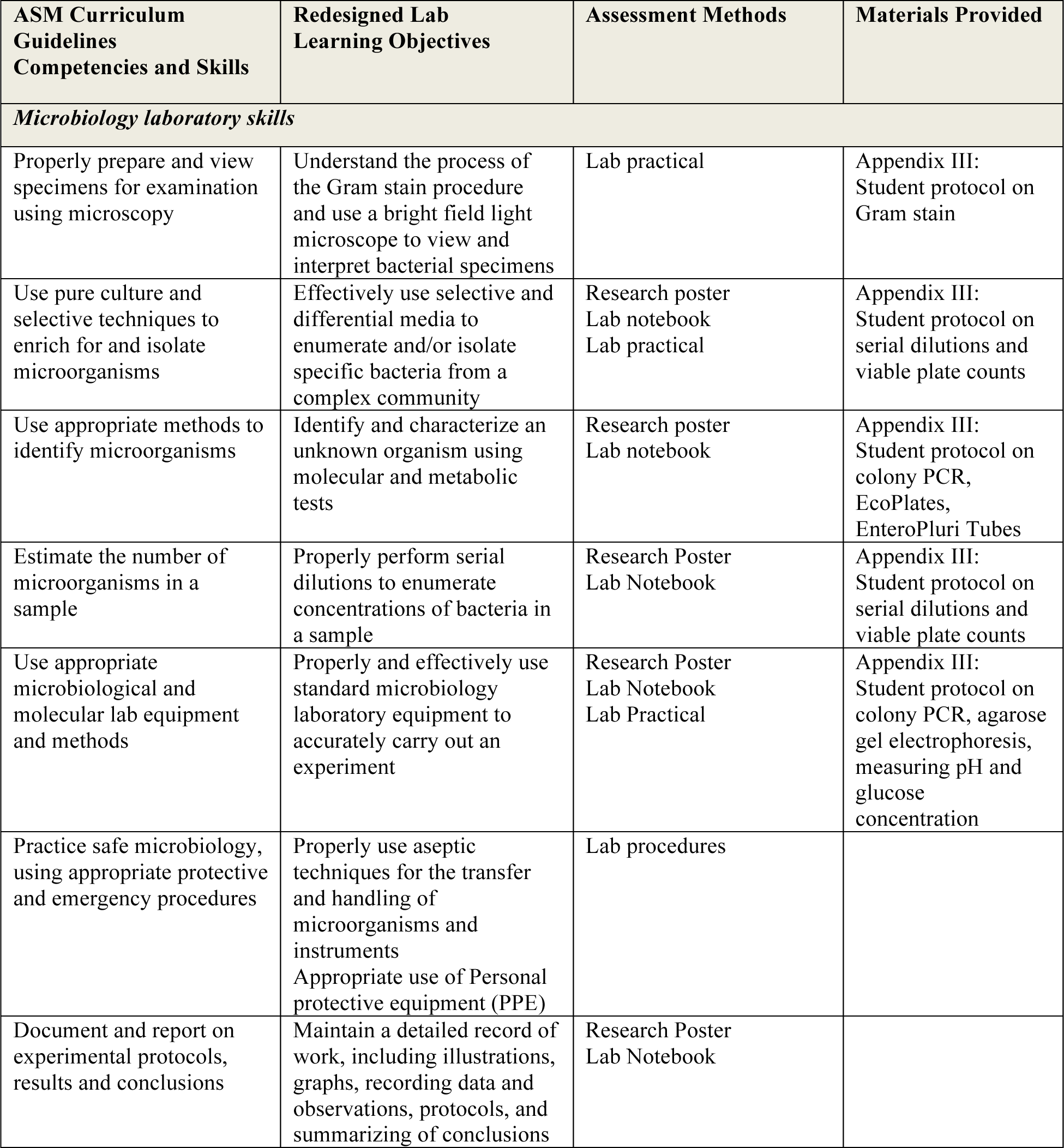
Mapping learning objectives to the ASM Curriculum Guidelines with corresponding methods of assessment and materials provided.

### Local Impact

Due to the nature of authentic experimental questions, the student-generated data will contribute to our understanding of microbial ecology in food fermentations and answer fundamental questions relevant to industry and the community on food safety and environmental impact. This project was in collaboration with the Mt. Olive Pickle Company, the largest privately held pickle company in the United States, headquartered in Mt. Olive, North Carolina. Knowledge of brining requirements may help inform their fermentation practices to reduce the amount of salts used in their brines, while producing a quality product and maintaining food safety.

## PROCEDURE

### Materials and Procedures

This lab course is divided into three separate but interconnected modules. Module 1 introduces core concepts and skills in microbiology including aseptic technique, media preparation, bacterial isolation, differential staining, microscopy, serial dilutions, bacterial enumeration, and food microbiology, that will be used throughout the inquiry-guided research project in Module 2. Module 3 is designed to teach and discuss applicable data analysis and data visualization techniques, including graphing and poster design. A weekly schedule of lab topics is available in Appendix I. Detailed weekly instructor preparation notes for the inquiry-guided project (Module 2) are included in Appendix II. Figure 1 illustrates the experimental timeline for Module 2, the inquiry-guided fermentation project.

**FIGURE 1.**
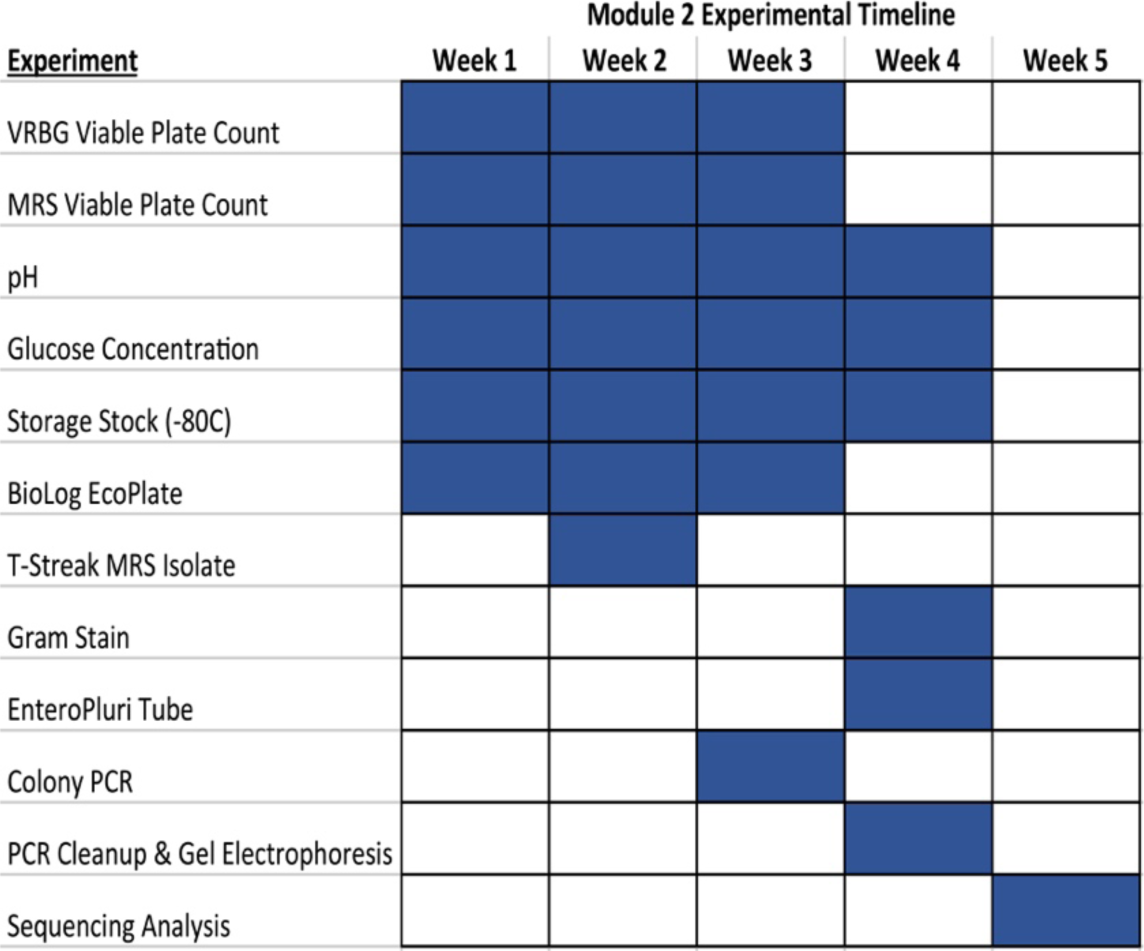
Weekly experimental timeline for Module 2: Inquiry Guided Research Project. Blue shaded boxes represent experiments performed that week.

The laboratory manual contains descriptive background information, detailed plans for each week, materials, lab protocols, media recipes, guidelines and rubrics for the laboratory notebook and research poster. It is important to note that our project is in collaboration with the Mt. Olive Pickle Company. Mt. Olive supplies all the cucumbers and fermentation jars for this course. Cucumbers, specific for pickling, can be purchased at local retail grocery stores, and fermentation jars can be reused 500 mL commercial pickle jars. The protocols for the cucumber fermentation in Module 2 include the Gram stain, cucumber fermentation jar set-up, serial dilutions, viable plate counts, measuring pH and glucose concentrations, Biolog EcoPlate inoculation, colony 16s rRNA PCR, PCR clean-up, agarose gel electrophoresis, and EnteroPluri tube test. Detailed student protocols are provided in Appendix III. For each protocol, the necessary materials and equipment are listed, followed by detailed step-by-step instructions. Student protocols are provided to students in the appendix section of the course lab manual, rather than embedded with the background information and goals for the weekly labs. This structure was specifically designed to encourage students to practice accessing protocols, which are repeated across multiple weeks, outside of a traditional in-text lab manual. Additionally, before the start of Module 2, an experimental design worksheet (Appendix IV) was given to students to help them prepare for their research project. The assignment requires students to state a research question, identify their selected experimental variable (NaCl concentration), develop a hypothesis based on their current knowledge, and graph predicted results of their fermentation experiment. Students are required to reference this assignment after the completion of the project as part of their discussion in their final poster presentation.

Module 2 is conducted over a period of 5-weeks. Briefly, groups of four students initiate their fermentation experiment by packing a 500 mL glass fermentation jar with cucumbers (~4-6 cucumbers) and their selected NaCl brine solution. This results in an approximate 1:1 ratio (weight:volume) of cucumber to brine. Each week, for 3-weeks, students analyze four specific parameters to assess the progress of their fermentations, including viable plate counts of *Enterobacteriaceae* (plated on VRBG agar) and *Lactobacillaceae* (plated on MRS agar), pH and glucose concentration of the brine solution, and community functional diversity (BioLog EcoPlates™). Groups work as a team to carry out all of the aforementioned experiments with one fermentation jar. It is advised that each week individual members of the group perform different experiments to gain experience with each method. Fermentation jars are incubated at room temperature (~22-24°C), and agar plates are incubated at 30°C. VRBG plates are incubated under atmospheric conditions, while MRS plates are incubated in anaerobic boxes (BD GasPak™ EZ anaerobe container system) in an effort to prevent the growth of yeast and molds. Cycloheximide can be used in the media plates in lieu of incubation in anaerobic boxes; however, Cycloheximide is toxic to mammalian cells, and use of the anaerobic box affords students an opportunity to work with anaerobic systems.

In addition to addressing community-wide assessments of their fermentations, students (working in pairs) identify and characterize a single MRS isolate from their fermentation experiment. Students select an isolated colony from an MRS plating experiment and perform a T-streak to obtain a pure culture. Students perform a colony polymerase chain reaction (PCR) to amplify the 16s rRNA gene. PCR amplification products are processed with a PCR clean-up kit and submitted for Sanger Sequencing (Eton Biosciences Inc.). In the following lab session, students are given their sequencing results as a text file, and are taught how to use BLASTn to help identify their isolate’s genus. Furthermore, students work to characterize the metabolic properties of their isolate using the EnteroPluri-Test (Becton Dickinson). The EnteroPluri-Test is not used for organism identification using the EnteroPluri-Codebook, but as a compact, and easy to use, method to analyze 15 different metabolic tests.

The final section of the lab (Module 3) is designed to teach students how to analyze and present all of the data collected over the semester in an electronic scientific research poster. Instruction focuses on data analysis, graphing techniques, and good practices for poster design. Although there is a dedicated Module for data analysis and visualization, these concepts are also incorporated into labs throughout the semester. Each week, students are encouraged to take their data and start the process of drafting a visual representation of the data, such as constructing graphs and tables. Example graphs were given to students to aid in their design (Appendix V).

### Suggestions for determining student learning

Throughout the semester, students are evaluated through several graded assessments, including three in-lab quizzes, laboratory notebook checks, lab procedures, a written final exam, and a final lab practical assessing core lab skills (T-streak, pipetting, Gram staining, and microscope usage). Furthermore, each student must design and create a scientific research poster presenting the results from their fermentation project. This poster is created in an electronic format (e.g. Microsoft PowerPoint) and does not need to be printed. Students upload a PDF document of their poster to a course management website (e.g. Moodle) for grading.

In this course redesign, we specifically focused on the final research poster as a major assessment of student learning. Although pre- and post-content quizzes offer insight into student learning gains, they are unable to easily and accurately assess a student’s global understanding of a research project. Assessment of a research poster can help indicate whether a student connects individual components of a project to the global objective and purpose; are they able to synthesize and conceptualize the results of their scientific project. For this poster assessment, students were given instructions in the lab manual and during lab lectures on what to include in their posters, a rubric outlining key components of the poster based on the ABRCMS Poster Judging Rubric^14^, online resources for creating effective scientific posters, and example posters from Graduate Student TAs in the NCSU Microbiology Graduate Program. The poster grading rubric is provided in Appendix VI.

### Sample Data

Examples of students’ graphs from different cucumber fermentations are provided in Appendix VII. Throughout their fermentation experiment, students enumerate populations of *Enterobacteriaceae* and *Lactobacillaceae* using viable plate counts, and track the pH and glucose concentration of the brine solution. A successful fermentation will result in the increase in concentration (CFU/mL) of *Lactobacillaceae* over time and a decrease in the concentration (CFU/mL) of *Enterobacteriaceae*. Glucose is the main sugar in cucumbers and over time will diffuse into the brine solution. This glucose serves as the primary carbon source fermented by microorganisms into lactic acid, resulting in a drop in pH. *Lactobacillaceae* are able to survive in this acidic environment, while members of the *Enterobacteriaceae* family are not. Based on student graphs, it is evident that students observed an increase in *Lactobacillaceae*, a decrease in *Enterobacteriaceae*, and a drop in pH over time, across all NaCl brine concentrations. The glucose concentration results were inconsistent, but typically yielded an increase after the first week and then a decrease, demonstrating utilization of glucose as a carbon and energy source. Use of all student data was approved by the NCSU Institutional Review Board (Protocol No. 12753).

### Safety Issues

Experiments were designed using BSL-2 lab procedures and the American Society for Microbiology Guidelines for Biosafety in Teaching Labs^15^. Experiments, were reviewed by the NCSU Institutional Biosafety Committee and documented in a Biological Use Authorization from NCSU Environmental Health and Safety (#2017-07-709). We chose to ferment cucumbers because they enable a safer fermentation process, whereas other vegetables, like okra, may result in incomplete fermentations, resulting in a biological safety risk. There is a potential that students may grow pathogenic microorganisms in their pickle brine solutions. To limit this, the minimum brine salt concentration students use is 2%, which we have demonstrated to be effective at inhibiting populations of *Enterobacteriaceae* (data no shown). Additionally, we monitor the pH of the fermentations to ensure that the pH remains acidic, below 4.6, an environmental condition that helps inhibit pathogenic microorganisms^12^. Moreover, students never handle or propagate isolates cultivated on VRBG selective media, as they may pose a health risk. MRS media is selective for *Lactobacillaceae*, and all propagated isolates are sequenced (16S rRNA gene sequencing) to ensure that they are non-pathogens. Moreover, students are required to wear personal protective equipment (disposable gloves, safety glasses, and lab coat) at all times. All biological and chemical waste is disposed of according to University policy.

## DISCUSSION

### Field Testing

The course redesign process occurred over four academic semesters with a team of two faculty members, a graduate teaching assistant (also the TA instructor for the lab), an undergraduate student who had previously taken the course, and a laboratory research technician. The first year of the redesign involved developing the inquiry-style of laboratory instruction, project selection, writing student learning outcomes, developing the laboratory manual, and evaluating the timing and feasibility of experiments. In year two, the redesigned course was piloted in two sections of MB 354 (the honors and majors section of General Microbiology Lab) in both Fall and Spring semesters. Piloting smaller sections (~15 students/section) enabled us to more closely observe student interactions, equipment usage, and space constraints. It also afforded us an opportunity to make small modifications to protocols, the weekly schedule, and instructional content to better serve student learning. For example, in the first semester piloting this redesign, we noticed deficiencies in the final project research posters. We realized that the instructions, rubric, and expectations of this project could have been more clearly stated throughout the semester. For the second semester pilot, we modified our rubric, instituted a poster peer-review process, incorporated discussions on poster requirements throughout the semester, and added a lab period on poster design, data analysis, and data visualization.

An example of an experimental issue was observed with the BioLog EcoPlates. These plates are a great tool to study functional diversity and metabolic potential of a community; however, the conditions in our brine solution consistently resulted in the negative control (water only) to change color, indicating metabolic activity, a false positive. Instead of removing this inconsistent assay, we kept it in the project to 1) teach the theory of functional diversity and the role of redox dyes in physiological screening, and 2) include an experiment where students need to understand the role of controls, learn to troubleshoot, and determine how to interpret experimental results when controls fail.

### Evidence of Student Learning

A major goal of this redesign was for students to participate in an authentic research experience while learning core microbiology skills and techniques. A key aspect of research is understanding that individual experiments are interconnected and come together to help answer a global research question. To determine if students were able to link individual experiments to the project as a whole, we assessed students’ final poster presentations. Specifically, we assessed if, and where, a core set of critical experiments were discussed in each student’s final poster (Figure 2). As a whole, many students were able to develop a descriptive title that described the research question, and included a descriptive and testable hypothesis within their poster introduction. Additionally, most students referenced their hypothesis in the discussion section.

**FIGURE 2.**
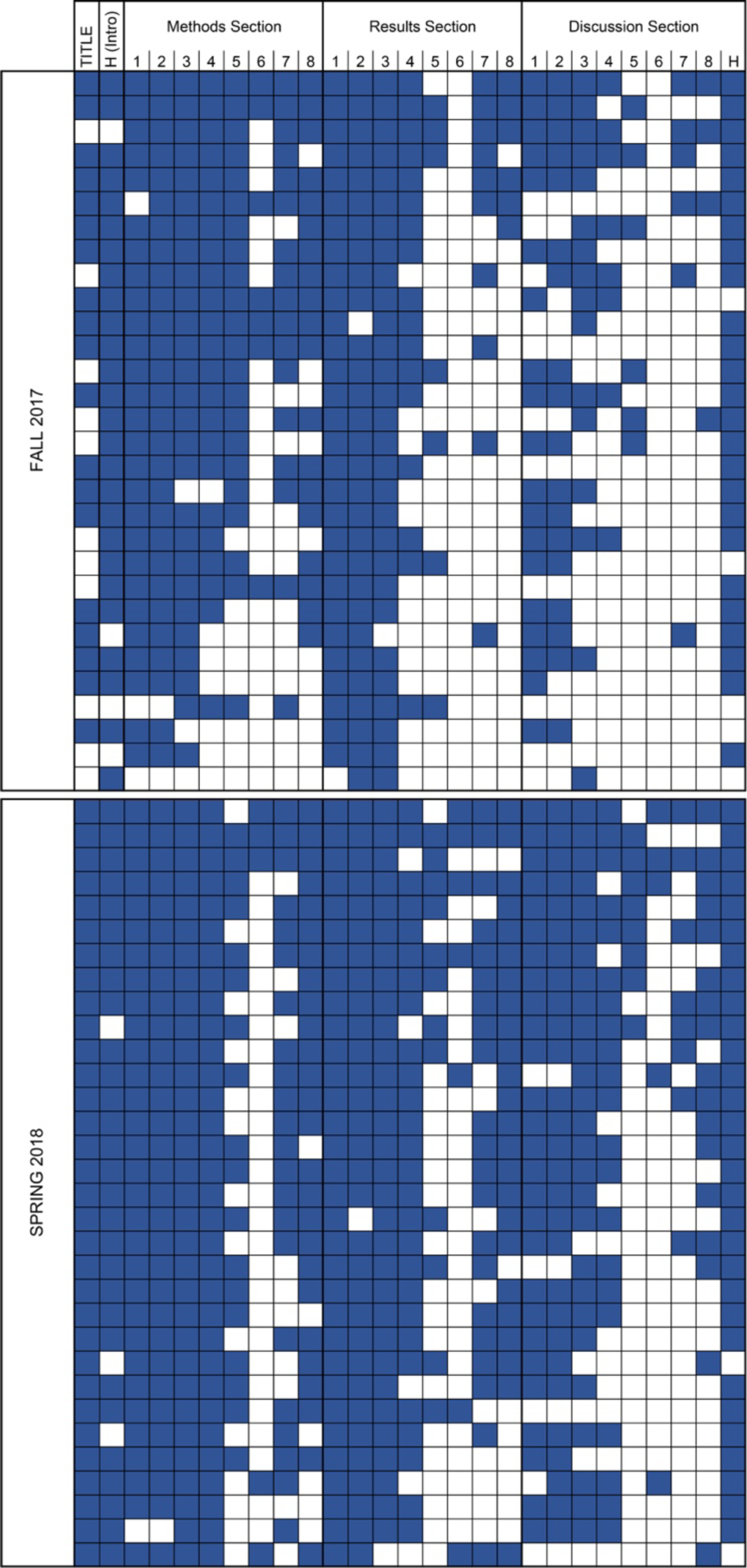
Assessment of student posters identifying the ability to connect individual experiments to the global research project. Student posters from two sections in Fall 2017 and two sections in Spring 2018 were aggregated and assessed to determine if students included specific project experiments and components throughout their final poster, including the Title, Methods, Results, and Discussion. Rows indicate individual student posters, and columns represent specific poster components (T: descriptive and scientifically valid title, H: inclusion of a testable hypothesis in the introduction and discussion, 1: VRGB plate counts, 2: MRS plate counts, 3: pH, 4: Glucose assay, 5: EcoPlate, 6: Gram stain, 7: EnteroPluri, 8: PCR/16S Sequencing. Blue shaded boxes indicate that the poster included the stated component, white boxes indicate that the component was absent from the poster.

When we assessed individual experiments, students were more likely to discuss each experiment used in the project in the methods section; however, they did not continue that discussion of each experiment, and its corresponding results, throughout the entire poster, including the results and discussion sections. Specifically, students were more likely to discuss experiments they repeated weekly throughout the research project, or experiments they perceived as “good,” specifically experiments with easily observable/interpretable results including CFU plate counts and pH measurements. Students were less likely to discuss experiments that were only conducted one time during the project, including 16S rRNA gene sequencing, Gram stain, and EnteroPluri Test, as well as experiments that produced sometimes inconclusive results, including the EcoPlate, and glucose concentration assay (Figure 2). The results from this assessment suggest that students may perceive experiments completed multiple times, with consistent and easily interpretable results, as more important than experiments conducted a single time. Moreover, students demonstrate difficulty connecting the experiments done on a single isolate to the overall research question.

Throughout the second semester piloting the redesigned lab, the importance of each experiment and the corresponding results to the global project were emphasized. Students were also given more direction about what to include in their research posters through a poster rubric (Appendix VI) and a peer review session. As a result, a more complete representation of their research projects was observed in students’ final posters (Figure 2). More students included information about their glucose assays, the EnteroPluri Test, and sequencing in their methods, results, and discussion sections. Students more frequently mentioned the EcoPlate in the methods section more, yet still did not state the results nor discuss them. Students again did not consistently include the Gram stain throughout their poster. It is possible students perceived this as a confirmatory test, as MRS isolates should be gram-positive, and did not differentiate it as a separate test. Finally, about half of the students failed to discuss their EnteroPluri test or sequencing results, even though they were more likely to include the EnteroPluri Test and sequencing in their methods and results. These results demonstrate that students continuously need to be analyzing their results, and thinking about where they fit into the global research project.

In an effort to determine if students were becoming proficient in basic microbiological lab skills and techniques in this redesigned course, we assessed a portion of the final lab practical exam. Specifically, we assessed students’ ability to perform a T-streak (obtaining individual/isolated colonies and maintaining a pure culture with proper aseptic technique), correctly classify an unknown bacterium as gram-positive or gram-negative using the Gram stain procedure, clearly focus a specimen under 100X oil immersion using a compound light microscope, and accurately use a micropipette (both a P1000 and P200). Although our sample size is small (30 students), overall, most students demonstrated proficiency in all assessed core lab skills (Figure 3). Some students had difficulty with the T-streak procedure, specifically obtaining individual/isolated colonies, as well as correctly performing the Gram stain procedure. Both of these skills are only performed twice throughout the semester. It is not surprising that 100% of students were proficient in pipetting, as this skill is a major component of the research project, and is repeated at minimum four times throughout the semester. These results lend some evidence that repetition may lead to increased proficiency.

**FIGURE 3.**
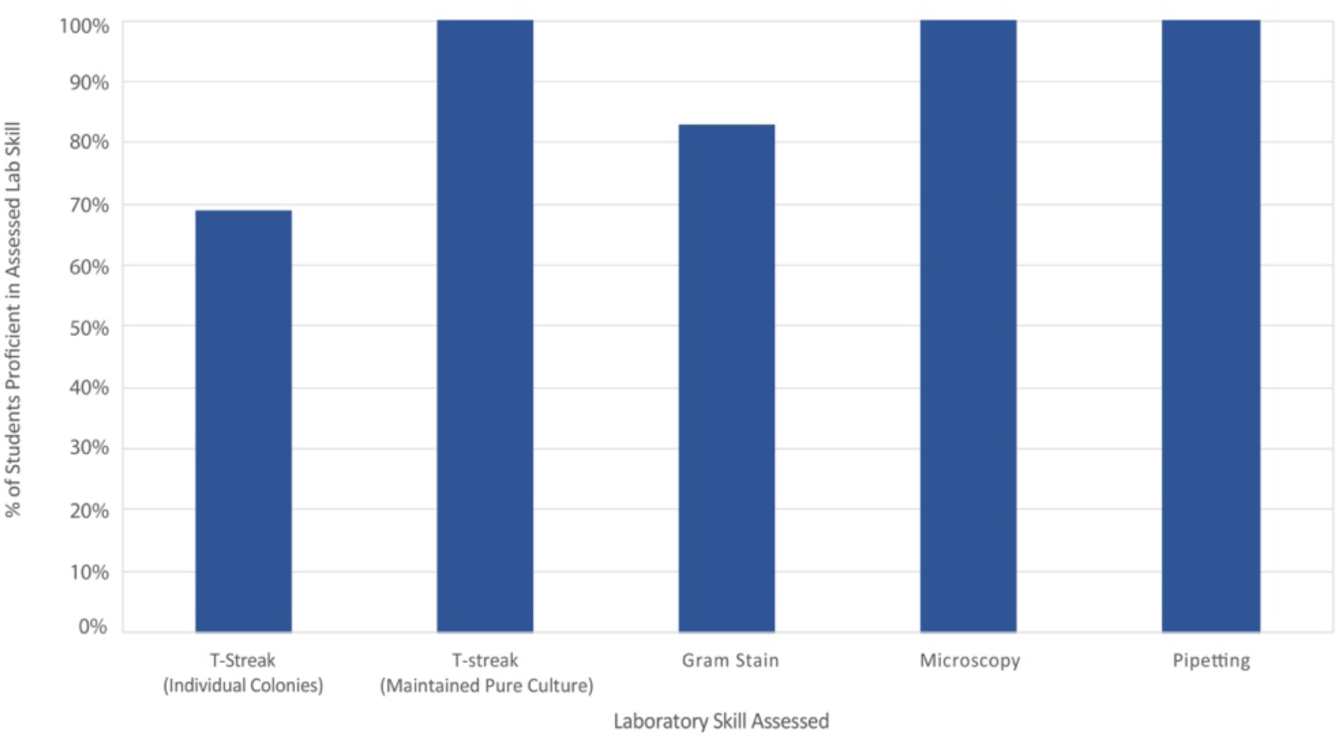
Assessment of student proficiency in a set of core skills and techniques during the hands-on practical portion of the final exam. Skills and techniques assessed include obtaining individual/isolated colonies and maintaining a pure culture with proper aseptic technique with a T-streak, correctly classify an unknown bacterium as gram-positive or gram-negative using the Gram stain procedure, clearly focusing a specimen under 100X oil immersion using a compound light microscope, and accurately using a micropipette (both a P1000 and P200). Bars represent the percent of students that were graded as proficient on each skill. Scores are aggregates from two different sections in Fall 2018, *N* = 30 students.

### Possible Modifications and Extensions

This introductory course is divided into three modules. Module 1 introduces basic microbiological techniques and experimental design during the first 4-weeks of course. Module 2 is the inquiry-guided research project that lasts 5-weeks, and Module 3 covers data analysis/visualization, and the lab practical, it runs 2-weeks. Dividing the course into three distinct modules will make it easier to incorporate different research projects into Module 2. Over time, the objectives of our project will change (i.e. different NaCl concentrations, different salts, different foods), with the potential of an entirely new research project. Additionally, the course could be modified for upper level microbiology courses. For example, a metagenomic analysis could be performed on the pickling brine throughout the experiment to assess global changes in bacterial populations. Additionally, if students already have pre-requisite microbiological skill knowledge, Module 1 could be modified allowing more time in Module 2 to expand the research project, including more replicates, adding additional metrics to assess, and/or modifying the research questions based on results gained in previous experiments.

## Acknowledgements

We would like to thank the Mt. Olive Pickle Company for their support in providing fermentation supplies and advice on project design, as well as the NCSU STEM Initiative Grant for partially funding this redesign, in addition to the support from the Biological Sciences Department at NCSU.

## SUPPLEMENTAL MATERIALS

**Appendix**

I. Weekly schedule of lab topics
II. Instructor Protocols
III. Student Protocols
IV. Student experimental design worksheet
V. Example graphs given to students
VI. Poster rubric
VII. Examples of student work

